# A vision prosthesis based on electrical stimulation of the primary visual cortex using epicortical microelectrodes

**DOI:** 10.1101/2020.12.15.422891

**Authors:** Denise Oswalt, P. Datta, N. Talbot, Z. Mirzadeh, Bradley Greger

## Abstract

Prostheses that can restore limited vision in the profoundly blind have been under investigation for several decades. Studies using epicortical macroelectrodes and intracortical microelectrodes have validated that electrical stimulation of primary visual cortical can serve as the basis for a vision prosthesis. However, neither of these approaches has resulted in a clinically viable vision prosthesis. Epicortical macroelectrodes required high levels of electrical current to evoke visual percepts, while intracortical microelectrodes faced challenges with longevity and stability. We hypothesized that epicortical microelectrodes could evoke visual percepts at lower currents than macroelectrodes and provide improved longevity and stability compared with intracortical microelectrodes. To test this hypotheses we implanted epicortical microelectrode arrays over the primary visual cortex of a nonhuman primate. Electrical stimulation via this array was used to evaluate the ability of epicortical microstimulation to evoke differentiable visual percepts. Visual percepts were evoked using the epicortical microelectrode array, and at electrical currents notably lower than those required to evoke visual percepts on macroelectrode arrays. The electrical current thresholds for evoking visual percepts on the epicortical microelectrode array were consistent across multiple array implants and over several months. Normal vision of light perception was not impaired by multiple array implants or chronic electrical stimulation, demonstrating that no gross visual deficit resulted from the experiments. We specifically demonstrate that epicortical microelectrode interfaces can serve as the basis for a vision prosthesis and more generally may provide an approach to evoking perception in multiple sensory modalities.

**One Sentence Summary:** Electrical stimulation of the brain via microelectrodes resting on the surface of primary visual cortex can evoke multiple differentiable visual percepts.

## Introduction

There are an estimated 39 million people in the world living with blindness[1]. Damage along the visual tract from either trauma or disease results in a variety of deficits with limited treatment options. Fortunately, over the last few decades, strides have been made in the realm of retinal prostheses with a slew of devices reaching preclinical and clinical testing, as well FDA and CE approval [2–8]. Prostheses interface epiretinally, subretinally or in the suprachorodal space and have been used to restore some limited functional sight in patients with retinitis pigmentosa (RP) and age-related macular degeneration (AMD). These approaches have been shown to increase acuity and improve quality of life [9], however patient populations are limited to those with functional retinal ganglion cells the degradation of which is common in many retinal diseases [10]. Patients with damage upstream from the retina, along the retino-geniculo-cortical tract are also excluded from a retinal approach. A neural prosthetic device providing stimulation of the visual cortex may restore functional visual to the profoundly blind when retinal stimulation in not a viable option.

The cortical approach for a vision prosthesis relies on the capacity for electrical stimulation of primary visual cortex (V1) to consistently evoke the perception of phosphenes. The foundation for this concept has been previously validated in studies using epicortical macroelectrodes with surface areas of .47 – 3 mm^2^ [11–14]. While these studies were able to evoke phosphene and in some cases simple patterns of light [15], certain fundamental limitations could not be overcome. The large surface area of the macroelectrodes resulted in high current thresholds, with some macroelectrodes requiring currents up to 8.1 mA [12]. Stimulation sometimes resulted in pain felt on the scalp or deep in the head. The sensations evoked were described as unnatural and in cases with high currents applied, multiple spots of light occurred, or the sensation lingered for minutes following cessation of stimulation [11, 12, 16]. The relatively large cortical areas being stimulated at high current levels also brought about concerns of evoking seizures [17], additionally limiting their capacity for concurrent multi-site stimulation.

For a visual prosthesis to be clinically relevant necessitates the use of high density electrode arrays that can consistently and safely induce phosphene patterns representative of a visual scene [18]. High-count microelectrode arrays, which interface with cortex at a physiologically relevant cortical column scale, could provide an effective platform for this purpose. Recent approaches in neural prosthetics have employed intracortical microelectrodes for their low current thresholds and high specificity in targeting cell populations [8, 15, 19–22]. Intracortical arrays consist of closely spaced penetrating electrodes and directly target deeper cortical layers [23]. Since intracortical arrays terminate near their targeted cell population they can provide more directed and localized stimulation at lower current levels compared to epicortical stimulation. Previous studies reported visual percepts evoked with intracortical simulation as low as 1.9-25μA [15, 21], however, stimulation of a few hundred micro-amps via several electrodes was routinely required, with thresholds increasing after several months of implantation before maintaining consistency [21, 22]. This decline is likely related to delamination of the electrical insulation on the intracortical arrays, glial encapsulation, and neural tissue damage [18, 24].

Greater tissue damage has been reported with intracortical than epicortical electrode arrays. Observed damage includes tissue dimpling, inflammation, foreign body response and encapsulation, decreased neural density near the electrode site, and persistent activation of microglia, potentially leading to neurotoxicity [25]. This effectively reduces the long-term viability of penetrating arrays and contributes to the increase in current required to evoke perception after chronic implantation. There is comparatively less documentation of the tissue response to chronically implanted epicortical arrays; however, because they do not penetrate the pial surface it is likely the foreign body response will be reduced [18]. Chronically implanted epiretinal microelectrode arrays remained functional after five years of implantation in human subjects [9]. Epicortical stimulation may similarly be capable of evoking photic responses after chronic *in vivo* implantation.

Array geometries pose additional considerations for tissue damage and surgical accessibility. Implanting intracortical arrays requires delicate manipulation as it pierces the parenchyma. Current intracortical arrays can be placed manually, but are more commonly placed with a pneumatic inserter [19, 20, 25]. Even with carful insertion, implantation cause neural tissue damage [25], which likely affects thresholds and tissue response. Conversely, epicortical arrays are more easily placed and repositioned, and cause minimal tissue damage [18, 26]. For all vision prosthesis systems, surgical implantation is complicated by the geometry of primary visual cortex. A large portion of the primary visual cortex lies within the sagittal fissure and calcarine sulcus. Epicortical arrays can be readily placed in the sagittal fissure. Sulcal implantation of both intracortical and epicortical arrays is very challenging, limiting the potential visual field representation. Access to these cortical surfaces could allow more complete representation of the parafoveal and peripheral visual field [18]. Although current thresholds for an epicortical microelectrode array may be higher than for an intracortical microelectrode array, using epicortical microelectrode arrays may provide reduced stimulation current thresholds relative to epicortical macroelectrodes and allow access to cortical structures less accessible to intracortical microelectrodes.

Experimental data suggests microelectrodes can simulate neural tissue at a distance 2-5 times that of the largest exposed electrode dimension [27]. Larger electrodes will be able to activate deeper cortical layers but require higher currents to maintain charge density, whereas smaller electrodes will have higher specificity but electrodes with too small of diameter may not be able to sufficiently activate the necessary layers. Studies using intracortical microelectrodes have often placed the electrodes in cortical layers two through four in part because of the low stimulation currents needed to evoke visual percepts [20, 28] and primarily because these layers could be useful for restoring vision as they include the termination of the geniculocortical projections (layer 4) and the origins of the cortico-cortical projections to higher level visual processing areas (layers 2/3) [29]. Epicortical microstimulation will likely be able to target the more superficial cortical layers 2/3. While studies have evaluated the efficacy and limitations of intracortical microsimulation [19-22, 30], there are limited reports regarding the capacity for efficacious treatment via epicortical microstimulation (ECMS). Given the potential benefits of a nonpenetrating, micro-scale approach, this study proposes a systematic verification and validation of the capacity for surface level microstimulation to evoke visual percepts with chronic implantation.

## Results

### Psychophysics: Behavioral responses to epicortical micro-stimulation

Epicortical micro-stimulation on individual 200μm diameter electrodes consistently evoked behavioral responses indicating perception of visual stimuli. Thresholds were collected on eleven of fourteen electrodes on implant 1, between 114 to 142 days post-implantation. A failure on the percutaneous connector (observed on day 144) limited further testing. For implant 2, thresholds were collected for thirteen of the fourteen 200μm electrodes on the array, as one electrode (D2) was non-functional. Initial thresholds were collected between days 16 to 34 post implant, and calculated from twelve of the thirteen functional electrodes, one electrode (B1) did not evoke any behavioral responses at the time of initial threshold collection. This electrode did begin evoking responses later in the study and is included in the chronic-stage averages, which were collected 244 to 262 days post implant. The initial average threshold across the arrays was 441μA for implant 1 and 324μA for implant 2. The chronic-stage average threshold for implant 2 was 316μA (Fig. 1A).

**Figure 1.**
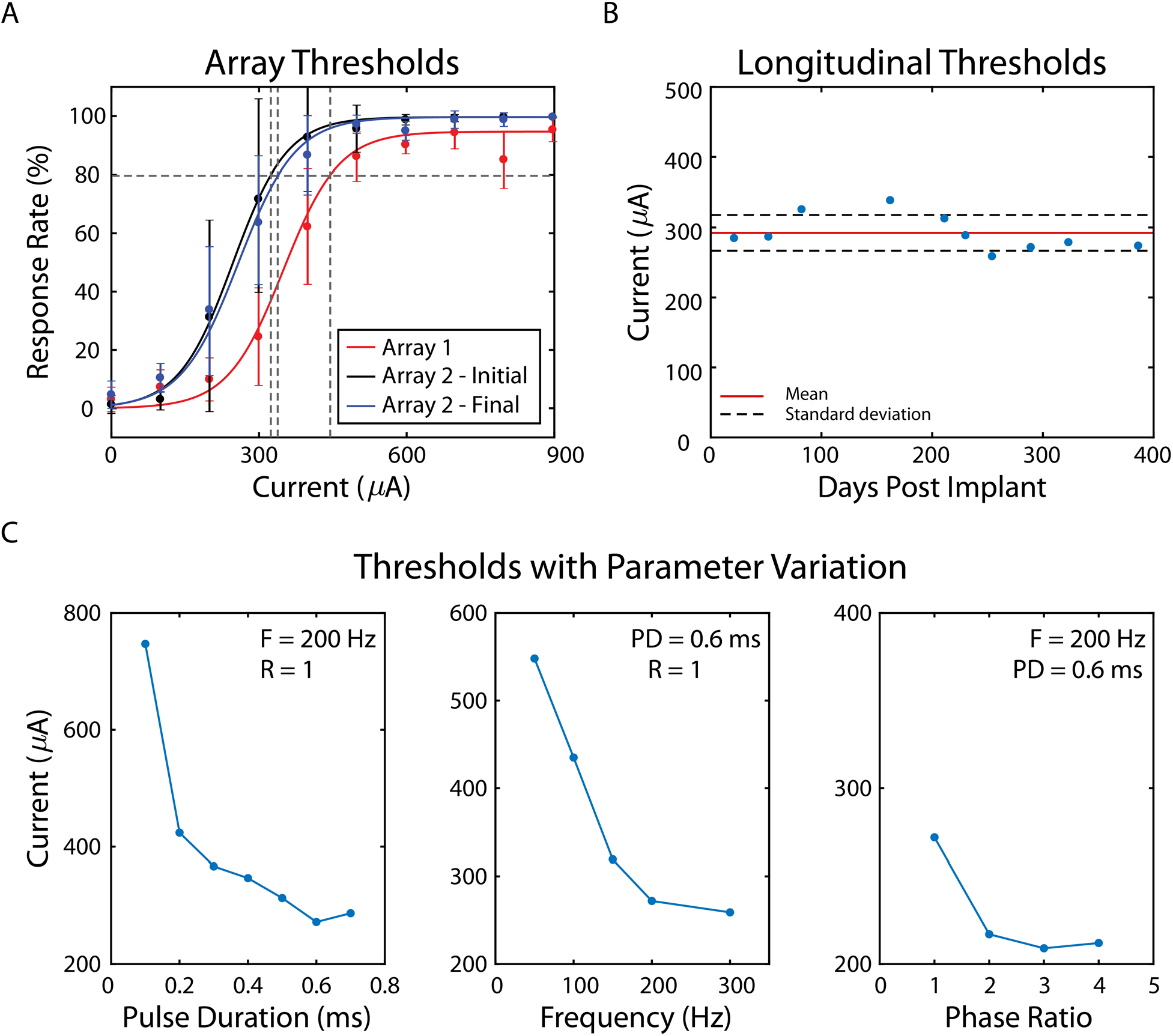
Electric stimulation threshold results. A. Average perception thresholds at 80% detection rate for 200μm electrodes on array 1 and array 2. Thresholds for perception were determined by fitting a cumulative Weibull function to data and setting 80% positive response probability as threshold, and are presented for 200μm electrodes. The red trace represents array thresholds for the first implant, black and blue are both thresholds collected for implant 2. Black indicates thresholds collected within the month following implantation, blue represents thresholds collected between 244 to 262 days post implant. B. Threshold stability over time for NHP1 implant 2. Data reported for electrode C2 on implant 2, with thresholds collected up to 386 days after the date of implant of the second array. The electrode had a mean threshold of 292.3±25.5μA. Mean indicated by red line with one standard deviation indicated by dashed lines. C. Effect of parameter variations on perception threshold. Left indicates the way in which threshold changes with pulse duration. Threshold decreases with pulse duration, before plateauing at longer pulse durations. Middle presents threshold as a response of pulse frequency. Increasing pulse frequency shows a decrease in threshold, and a plateau at higher frequency values. Right shows threshold response to phase asymmetry. Perception threshold decreases with a longer charge reversal duration.

Current stimulation thresholds collected on a single microelectrode (C2) of implant 2 showed consistent perception thresholds across a 12.8-month period, with an average of 292±25.5μA (Fig. 1B). The initial threshold collected 21 days post implant was 285μA; the threshold collected 386 days post implant was 274μA. The lowest threshold collected for this electrode was collected on day 254 and was 259μA. the highest threshold collected was 339μA and collected at day 162.

Individual micro-stimulation parameters were systematically varied to observe the effect of phase duration, pulse frequency, and phase asymmetry on the current thresholds required to evoke visual percepts (Fig. 1C). Threshold was observed to decrease with longer pulse durations, increased pulse frequency, and phase asymmetry. For phase durations of 0.1ms to 0.6ms, threshold decreased from 747μA to 272μA respectively. Increasing pulse frequency from 50Hz to 300Hz decreased threshold from 548μA to 259μA. Slowing the rate of charge reversal with charge-balanced phase asymmetry improved thresholds from 272μA for the symmetric case to 209μA for an asymmetric ratio of 3.

### Safety: Electrical charge and impact on visual function

Thresholds collected for pulse width durations from 0.1ms to 0.8ms were used to create perception level strength and charge duration curves for stimulation trains pulsed at 300Hz for 300ms on one electrode (C2) (Fig. 2). Data were collected from 254 to 264 days post implant. Rheobase was determined to be 502μA, and chronaixe at 0.185ms. Minimum charge was 82nC/ph.

**Figure 2.**
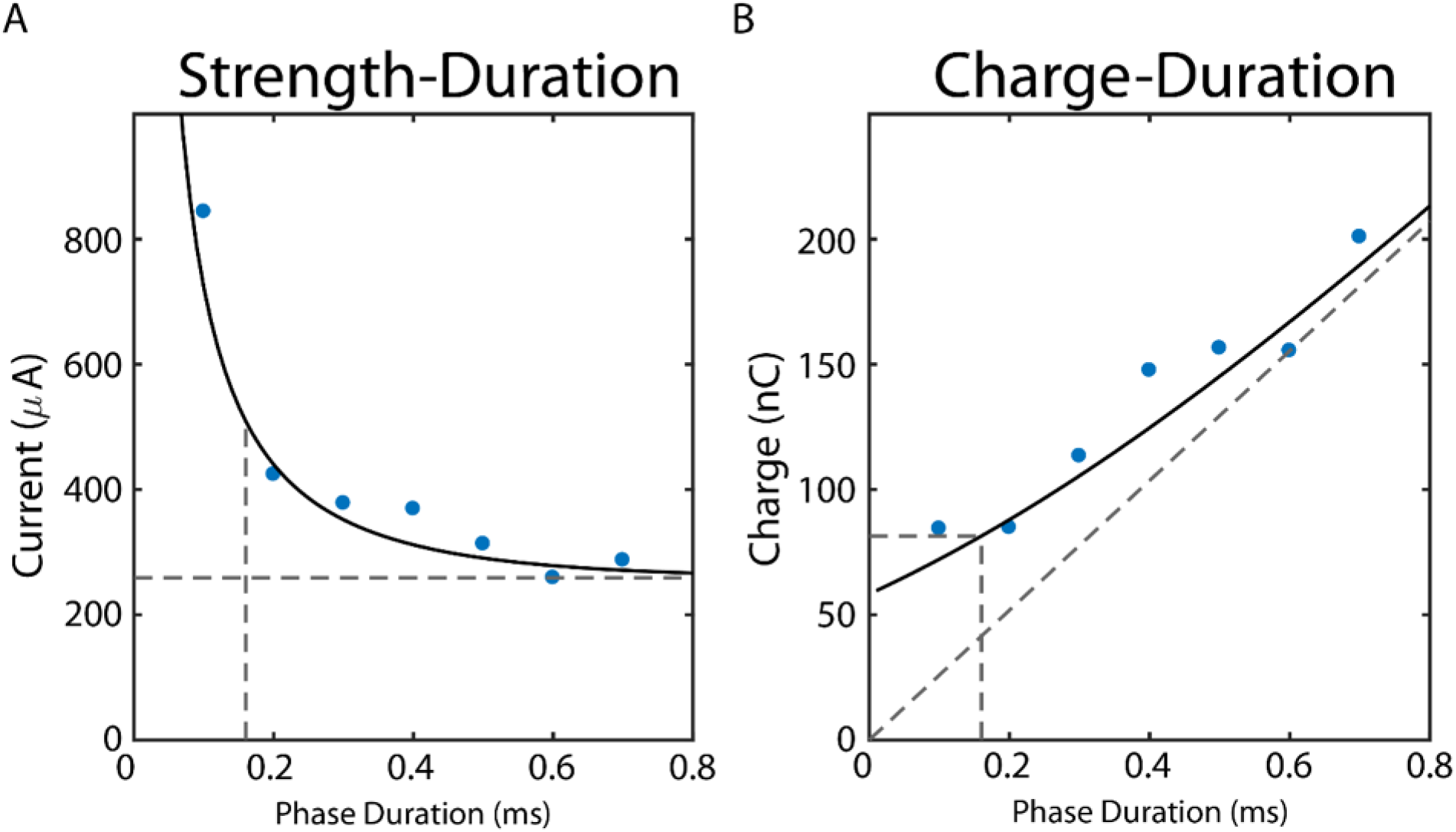
Strength and charge duration curves collected based on 80% perception thresholds. A. Current thresholds for each tested phase duration. Solid black line is the current threshold calculated in accordance to its relationship to measure rheobase and an assumed time constant. B. Charge duration curve at threshold values with calculated minimum current. Each trial conducted consisted of 300 Hz pulse frequency and 300ms train duration.

The ability of the nonhuman primate to detect photic stimuli placed within visual field of the area of primary visual cortex underlying the epicortical microelectrode array was used as a measure of cortical tissue function. An increase in the threshold of photic stimulation needed for detection would be an indication of dysfunctional tissue, and in the extreme would represent a visual lacuna. Luminance values of 10-45cd/cm^2^ and 60cd/cm^2^ observed no decline in photic responsiveness from the pre-to post-study condition (Fig. 3A). Stimuli presented at 45-50cd/cm^2^ did observe a decline in average responsiveness but deviations were not considered statistically significant (t-test, p>0.01).

**Figure 3.**
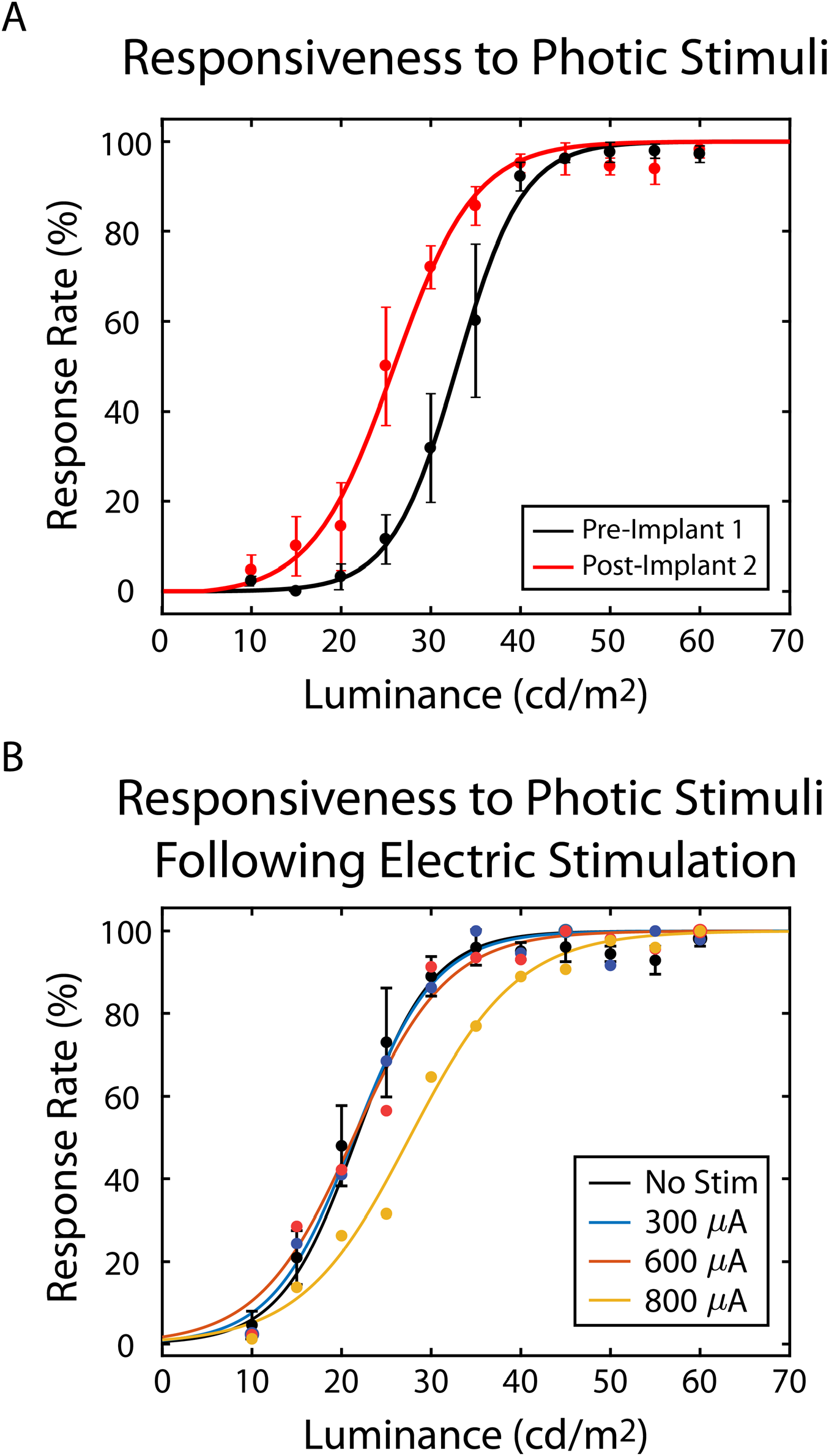
Responsiveness to photic stimuli. A. Photic psychophysics pre- and post-study for NHP1. Responsiveness to photic stimuli with varied luminance was observed to increase for luminance from 10-45cd/cm^2^ and 60 cd/cm^2^. Responsiveness decreased for luminance values from 50-55cd/cm^2^. B. Psychophysics curves for photic stimulation trials in the 500ms following cessation of electric stimulation, on an electrode with 293±25μA threshold. The black curve indicates the psychophysical responsiveness to photic stimuli without electric stimulation applied during that session. The blue curve represents photic stimulation in the trial window following termination of stimulation applied at a 300μA. The red curve indicates responsiveness to photic stimuli following stimulation at 600μA. The yellow cure indicates photic responsiveness following application of an electric stimulus of 800μA.

Responsiveness to photic stimuli within 400ms following a 300ms pulse train of electric stimulation was used to characterize the temporally lingering effects of stimulation on endogenous visual processing. For an electrode with threshold of 293±25μA, psychophysics indicates no significant divergence in responsiveness to photic stimuli at each luminance value tested when electrical stimulation is applied at 300μA or 600μA (t-test, p>0.01). Results indicated significant deviations in responsiveness for luminance values 20-45cd/cm^2^ (p<0.01) (Fig. 3B). Psychometric curves for the 600μA and 800μA condition shift their respective psychometric curves to the right, increasing the photic threshold for perception. With the no stimulation condition, the photic threshold was determined to be 27.4cd/cm^2^. The condition following 300μA stimulation returned a photic threshold of 27.5cd/cm^2^. For 600μA, this is increased to 28.6cd/cm^2^ and to 35.8cd/cm^2^ for the 800μA condition.

### Acuity factors: Two-point discrimination

A two-point discrimination task was used to assess if spatially differentiable visual percepts could be evoked by simultaneous stimulation of two electrodes. The behavioral responses of NHP1 during this task indicated consistent perception of spatially differentiable visual percepts. NHP1 reported that differentiable visual percepts were observed in 90% of trials in response to simultaneous stimulation on the closest neighboring electrodes, 2.08mm center-to-center distance. Larger distances between stimulating sites continued to yield responses indicating separate percepts (Figure 4). The analysis consists of 31 electrode pairings and 5009 stimulation trials. The behavioral responses of NHP1, during “catch” trials in which electrical stimulation was applied only to a single electrode (distance = 0mm), indicated differentiable visual percepts in only 11% of trials.

**Figure 4.**
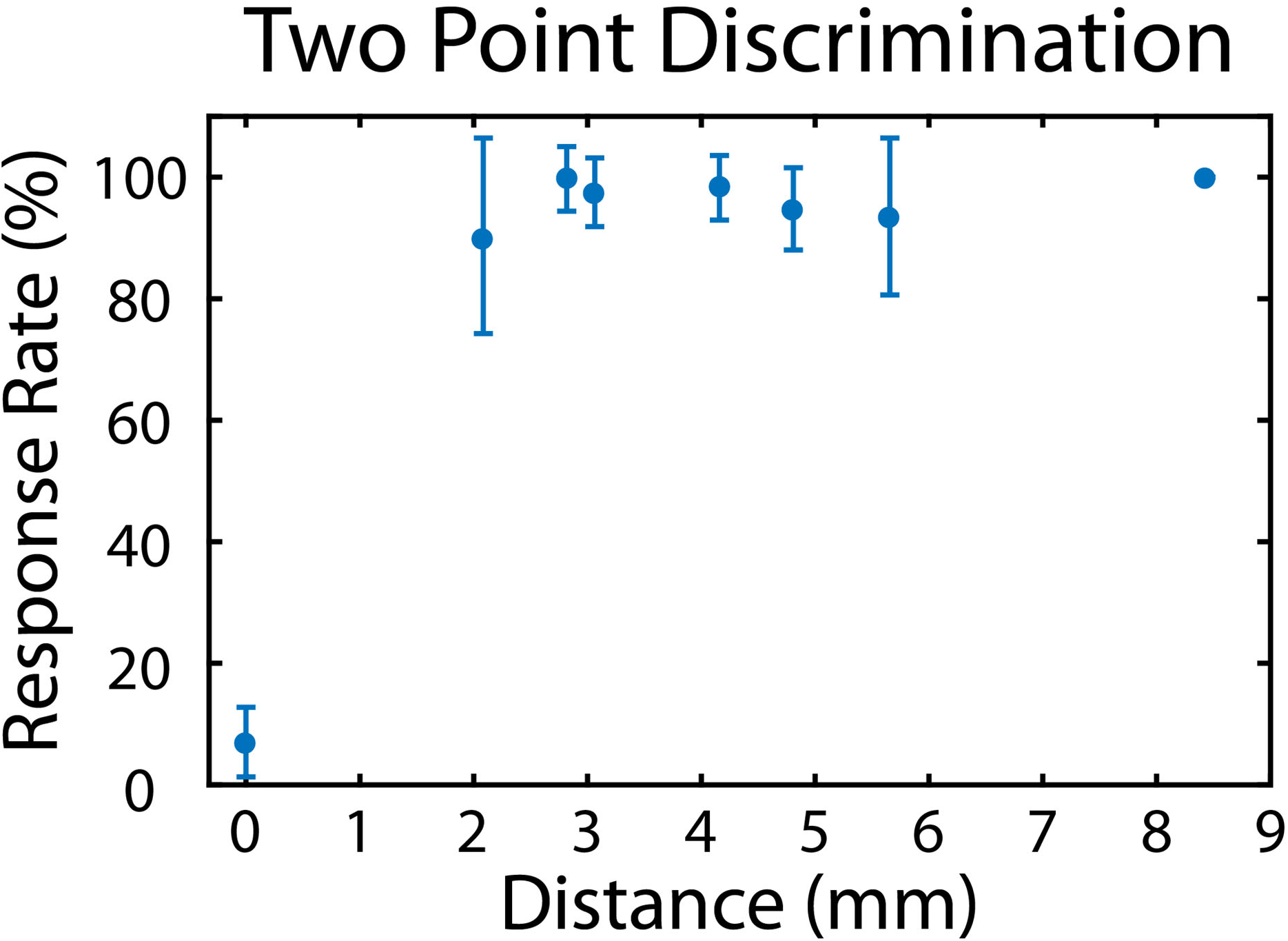
Two-point discrimination psychophysics. NHP1 behavioral responses to simultaneous stimulation on two electrodes with varying inter-electrode distances. Response rate reflects rate behavioral responses indicating perception of differentiable visual percepts. A distance of zero millimeters indicated stimulation at a single electrode site and was used as control. NHP1 behavioral responses to stimulation on two electrodes indicated perception of separate percepts at each distance tested. Nearest electrode pairs were separated by 2.08 mm, center-to-center.

The effective range of stimulation and percept size were predicted based on the array-average threshold (316μA). With a K of 675μA/mm, threshold stimulation returned a radius of 0.66mm. In relation to striate cortex geometry, this translates to a depth of activation through layers II/III and some activation of IVa and a width on the scale of a hyper column (Fig. 5A). Phosphene size is predicted to remain under 1° in diameter below 8° of eccentricity for a conservative estimation (upper bounds), with lower bounds predicting phosphene sizes under 0.5° for eccentricities below 14° (Fig. 5B).

**Figure 5.**
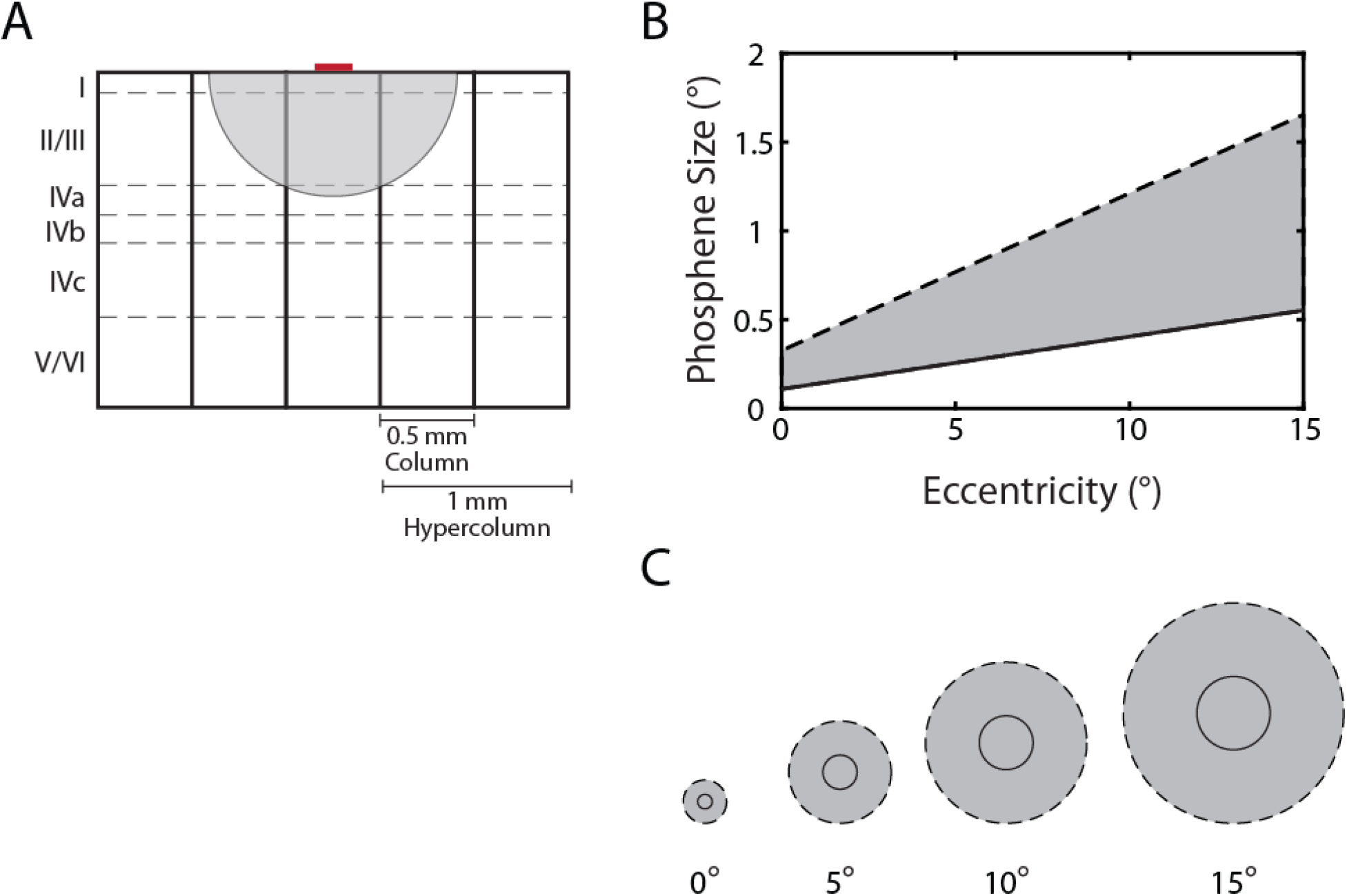
Estimated stimulation radius and percept size. A. The radius of activation at threshold stimulation. The red region indicated the 200μm electrode on the surface of cortex. The grey shaded region indicates the range of tissue activated by a current of 316μA, given a K = 675μA/mm^2^, in relation to layers in striate cortex. At threshold, stimulation reaches layers II/III and the upper portion of Iva. B. The range of predicted phosphene sizes at threshold stimulation for indicated eccentricity values. The solid black line reflects the lower bounds, the dashed line signifies the upper size bounds. C. Comparison of size values for stimulation at increasing eccentricity. The out circle bounded by the dashed line indicates the upper size bound, the solid black line indicates the lower bound for phosphene size.

Spatial patterns in electrical current stimulation thresholds were observed, with thresholds tending to increase with eccentricity along visual field and remining consistent at similar values of eccentricity (Fig. 6). This was observed for both implant 1 and implant 2. With implant 1, electrodes placed at increasing eccentricities, B2 - B4, observed thresholds of 290μA, 373μA, and 445μA. Electrodes with similar eccentricities, B3 and D3, returned similar threshold values 373μA and 375μA. For implant 2, B2 - B4 were positioned with increasing eccentricities and reported thresholds of 151μA, 190μA, and 240μA. Conversely, C4 and D3, at similar ranges of eccentricity, reported threshold of 370μA and 371μA respectively. This trend did not persist following longterm stimulation on implant 2, with most electrode thresholds drifting closer to the mean value. The lowest threshold observed on the array immediately following implantation was 151μA on B2, the highest was 495μA on C1. After eight months, the lowest threshold was 186μA on C3, and the highest was 475μA on A2.

**Figure 6.**
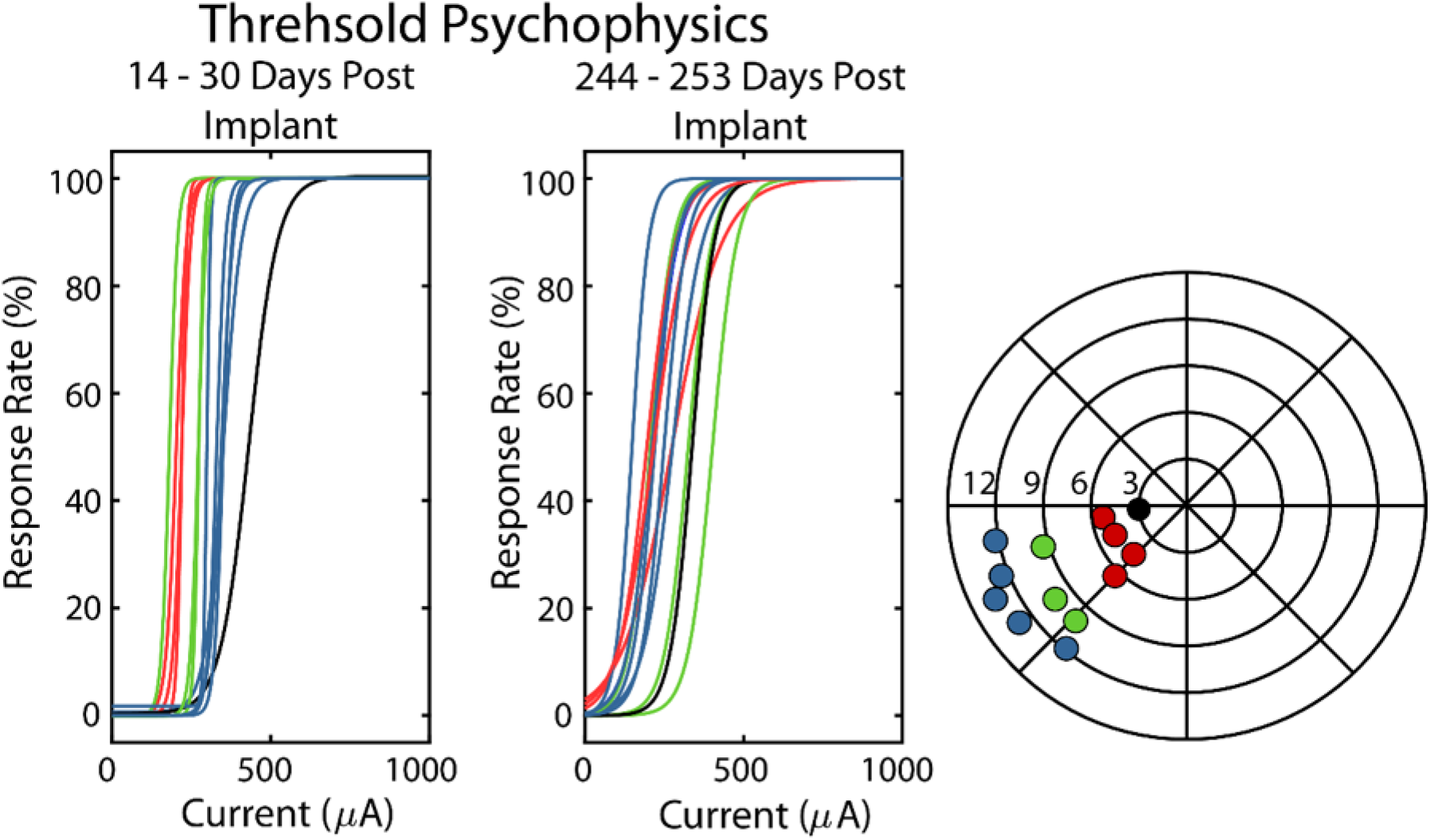
Spatially varying trends for perception thresholds. Right most polar plot indicates color groupings of electrodes at three similar eccentricities, and one electrode Left plot shows a general trend of increasing threshold with increasing eccentricity. Middle plot shows a dissipation of initial trend after several months of stimulations. Right most polar plot indicates spatial mapping and color code for eccentricity groupings determined from receptive fields.

## Discussion

ECMS was able to consistently evoke visual percepts in an NHP model. This constancy extended to implants over the same area of cortex and a combined implant time over a year. Thresholds were significantly lower than the several milliamperes required to evoke perceptual responses with epicortical macroelectrodes and counters the claim that electrode diameter does not have an impact on perception thresholds [12]. This claim however, was validated only with electrodes over 1mm in diameter. Epicortical electrodes of this size are thought to operate on a different scale than submillimeter-diameter microelectrodes [27, 31, 32]. Microelectrodes have a significantly more localized interface with tissue. The charge necessary to effect change on a smaller volume of tissue is expected. The extent of how localized that volume is a notable factor for epicortical electrodes for main reasons, the first being whether it can sufficiently active enough tissue in layers deep enough to evoke perception, the second relating to neighboring stimulation cites and minimum perceptible difference. A previous study was able to evoke visual percepts with electric stimulation in layer 1 of rhesus macaque striate cortex with 127μm wire electrodes [20], and another group evoked visual sensations with 500μm non-penetrating electrodes []. We were able to consistently evoked percepts with ECMS stimulation on 200μm electrodes. With feasibility demonstrated, additional factors involved in consistency, safety, and resolution can be evaluated.

For electrical stimulation-based prosthetics it is imperative to maintain low stimulation thresholds to avoid stimulation-induced tissue damage; it also factors into the effective visual resolution of a finalized prosthetic device. Lower current amplitudes activate smaller volumes of cortex, allowing neighboring stimulation sites to be closer without interfering with each other. Higher thresholds translate to larger volumes of tissue being activated, increasing the minimum distance at which two neighboring electrodes can evoke distinct sensations. In this study, the subject consistently reported two distinct percepts from concurrent stimulation on electrodes with a center to center spacing of 2 mm. The absolute minimum distance for 200μm electrodes may be smaller, but this remains to be tested in electrode arrays with tighter inter-electrode spacing. This distance was evaluated with an applied current of 600μA, supra-threshold for all electrodes. Since current spread has a smaller radius with lower amplitudes of applied current, it is likely that two-point discrimination can be maintained with closer inter-electrode spacing if conducted at threshold. The data presented was averaged across all electrode pairings tested, including those with electrodes placed at greater eccentricity. The NHP consistently responded in a manner indicating perception of two visual sensation for all pairings conducted. Considering the pairings

Two-point discrimination at 2mm center-to-center spacing was consistent across the array, including more eccentric electrode pairings. and held true regardless of eccentricity. Differentiation will likely be influenced by this current, with greater perceived separability likely with lower applied currents.

For a prosthetic device to be worth the risk of implantation requires it to provide therapeutic benefits consistently over time; minimizing tissue damage can increase the functional longevity of a device. Given the physics and placement of epicortical electrodes, ECMS tends to have higher baseline thresholds and larger volumes of activation compared to ICMS. However, they potentially offer a more stable system with better long-term viability in vivo. Intracortical studies observed an increase in threshold after several months of implantation [21, 22]. Epicortical arrays may maintain more consistent thresholds over time since they evoke a comparatively minimal tissue response. Similar technologies, such epiretinal stimulation has shown considerable consistency for over five years [9], and other implantable systems that employ long term subdural stimulation, such as the NeuroPace RMS System (NeuroPace, Mountain View, CA, USA) have also demonstrated therapeutic responsiveness to stimulation for over two years[33, 34]. This study evaluated ECMS in NHP V1, showing consistent perception thresholds with two arrays and longitudinally across several months. Additionally, the array was safely explanted after seven months of implant, with a new array placed over the same area. This is not typically feasible with intracortical arrays due to the tissue dimpling caused by their placement and the general fragility that hinders explanation. With the epicortical approach detailed herein, response thresholds did not increase, nor was photic perception impaired. Provided this stability continues, an epicortical approach may provide a more consistent platform for a prosthetic system.

A review of therapeutic electrical stimulation concluded damage thresholds for microelectrodes are more dependent on charge per phase rather than charge density as with macro electrodes, with pulse frequency also being an important factor [27]. While strict safety limits have yet to be determined for microelectrodes, microelectrodes have been shown to safely provide current densities exceeding the 30μC/cm^2^ level approved by the FDA [27, 31, 32]. This safety range was tested specifically for microelectrodes with a geometric surface area (GSA) of 0.00020.002mm, whereas the traditional Shannon model of damage was derived from electrodes with a GSA in the range of 1-50mm^2^. Microelectrodes, particularly surface microelectrodes, like the ones used in this study, typically have GSAs 0.03-0.07mm^2^ and occupy a space in-between these two metrics. As these electrodes do not fall into the GSA ranges for which the predictive damage models were established, specific testing of the acceptable charge density and charge per phase limits for these larger microelectrodes will likely be necessary. Maximum charge density and charge per phase applied here were 1.1mC/cm^2^ and 540nC/ph respectively, far above threshold values of 0.177-0.35mC/cm^2^ and 87-172nC/ph. With this level of current injection, no functional deficits were detected. Pre-and post-study responsiveness to photic stimuli used as a metric of visual function did not indicate acquired defects to natural perception nor lasting disruption of local visual processing. NHP1 did show an increase in responsiveness to lower luminance values, which is likely a training effect, rather than a sensitization caused by electrical stimulation of cortex.

The cortical magnification factor (CMF) in visual cortex means more cortical area is allocated to visual field space located near the foveal representation; the cortical area allocated per visual degree decreases with eccentricity. With a linear simplified CMF, this means evoked phosphenes are predicted to increase in size with eccentricity for a set current []. Additionally, the relationship between applied current and the radius of activated tissue dictates that increasing current also increases the effective radius (Eqn. 3). In this study stimulation thresholds across the array initially followed a general trend of increasing with eccentricity. Based on the mathematical relationships between applied current, activated cortex, and phosphene size, the threshold trend means the electrodes placed over more peripheral representations likely evoked much larger percepts at the time of initial threshold collection. For example, an electrode at 4° with a determined threshold of 151μA will evoke a phosphene of 0.16-0.48° in diameter, whereas an electrode placed at 11° with a threshold of 240μ is predicted to evoke a phosphene of 0.39-1.17°. A recent study applying epicortical stimulation on large ECoG electrodes in humans reported a saturation in phosphene size over a certain level of applied current [36]. Provided this phenomenon is consistent with micro-stimulation, size discrepancies may be less dramatic between percepts evoked in the foveal and more peripheral visual field representations. For a vision prosthesis, maintaining consistent percept size would be ideal to improve consistency and limit image distortion when evoking complex imagery with stimulation. Furthermore, the spatial trend for thresholds, while clear with initial array thresholds, was not observed following across the array after chronic stimulation. Thresholds for electrodes in more peripheral representations tended to decrease, while more foveal electrodes tended to maintain similar thresholds or slightly increase. As a result, thresholds across the array were generally more consistent. The initial trend may have been attention related as well as represent an adaptive learning effect. Previous studies in primates have shown decreases in response thresholds to visual cortex stimulation with prolonged training during forced-choice detection tasks [37]. This would account for reductions in thresholds observed from implant 1 to implant 2, as well as longitudinally with electrodes on the second array.

The main advantage in targeting the primary visual cortex for visual prostheses is the cortical magnification factor (CMF), which relates a distance across a sheet of neurons, representing a given angle visual field space [38, 39]. The upper range of CMF for foveal representations in humans and nonhuman primates has been reported to 15-20 mm/°, compared to a magnification factor of approximately 0.3 mm/° in the retina [11, 38, 40–43]. For visual prosthetics, this translates to a higher resolution being attainable when targeting V1. The values for CMF provide some insight into what level of resolution or artificial acuity that may be attainable with the approach detailed in this study. The results obtained suggest clearly separable percepts can be evoked at a distance of 2 mm, suggesting sub visual degree resolution is attainable with ECMS. Predictive calculations further support this, with upper limits for prediction indicating sub-visual degree phosphenes for percepts evoked within 8° of center, and lower limits indicating phosphenes smaller than half a visual degree are attainable within 14° of center.

## Materials and Methods

### Grid design

An epicortical array featuring concentric rings was designed and produced in tandem with Second Sight Medical Products (Sylmar, CA 91342, USA). The array consists of 46 electrodes. The center electrode for each grouping has an exposed area of 200μm. Each concentric ring is composed of these 200μm electrode pads electrically tied together. The concentric rings are composed of 6, 12, or 18 electrode pads, creating effective diameters of 750μm, 1500μm, and 2100μm. The array has a length of 13.84 mm and height of 9 mm. Groupings have a spacing of 2.12mm from ring center to center. Four rings with increasing eccentricity across the midline of the array contain each of the four electrode diameters; all remaining groupings contain the center electrode, the 6-ring, and the 12-ring (Figure 3). The array was potted to a titanium base with ZIF clip connector. The pedestal base is used as ground.

**Figure 9.**
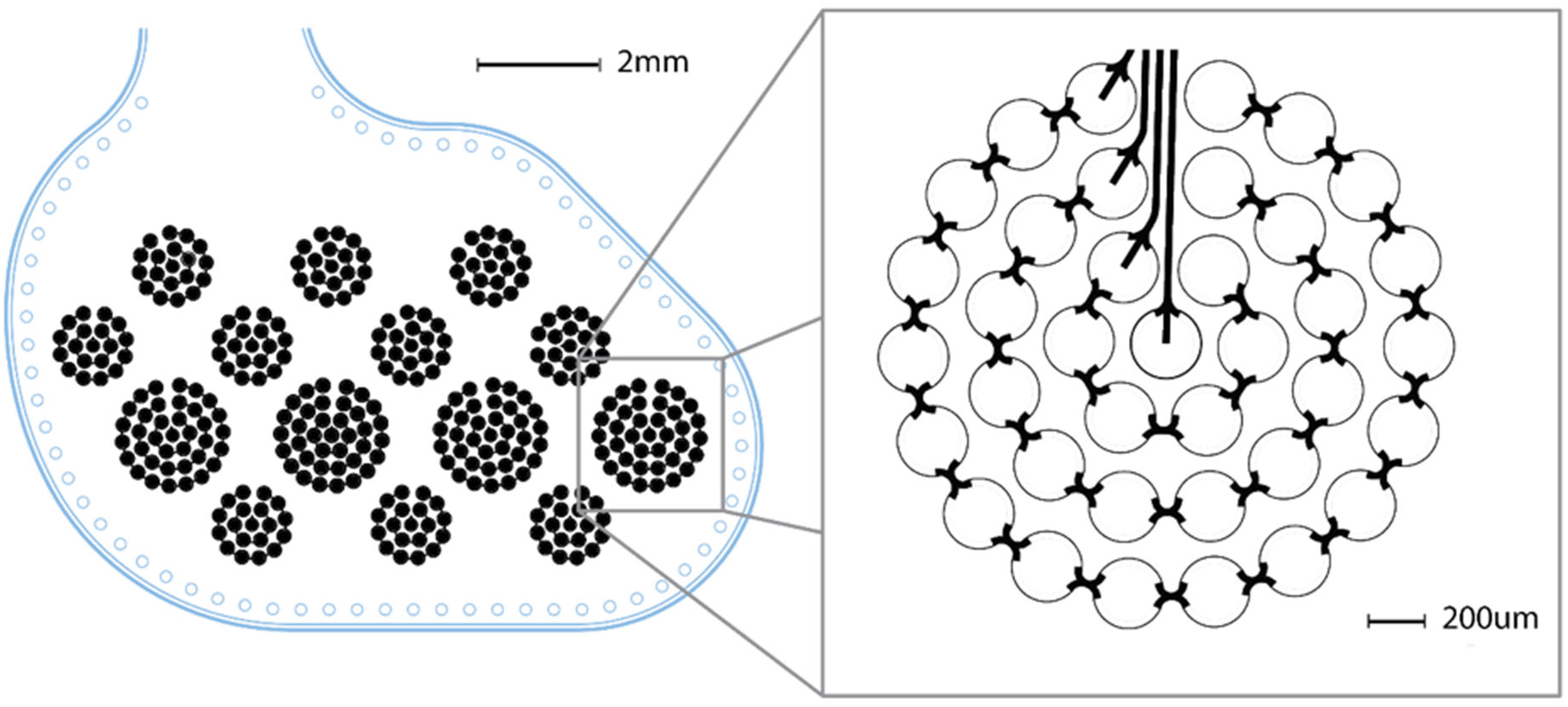
Custom array design with concentric rings. Array allows for probing thresholds and percept characteristics while accounting for variations with eccentricity and maximizing spatial landscape of the array. Rings made up of 1, 6-, 12-, 18-disk groupings and provide electrode diameters of 200, 800 1400 and 2000μm.

### Subjects and surgery

One adult male rhesus macaques (Macaca Mulatta) will be used in this study, NHP1. NHP1 was implanted with the epicortical array described above. The array was placed in the sagittal fissure facing the right hemisphere, with the caudal-most point of the array placed in close approximation to the pole, superior to the calcarine fissure. Human protocols were followed during surgery and post-operative care.

### Threshold detection task

This task is used to determine the luminance threshold for detection of photic stimuli as well as current thresholds for perception of electric stimuli. The NHP is placed in a primate chair inside of a dark chamber, facing a CRT monitor. The subject is required to place each hand on a capacitance switch, one to the left and the other to the right. A small red dot (.1X.1 visual degrees) at the center of the CRT screen was used for a fixation point. Once the NHP directs its gaze within one degree of the center point, an auditory tone signals the start of the trial. The NHP must maintain eye fixation for 400ms before a stimulus, either photic or electric, is presented. Electric stimuli are described below. Photic stimuli were round, monochromoatic Gaussian shapes with a diameter of one visual degree. Luminance levels are varied between 10 and 60 cd/m^2^ in 5 cd/m^2^ increments and presented on a screen with background luminance of 10 cd/m^2^. Photic stimuli are presented for 300ms. Following a hold duration of an additional 300ms, a second, unique auditory tone cues the NHP to respond. The NHP responds by removing its right hand to indicate that it did not perceive, or its left hand to indicate that it did perceive a stimulus. Photic threshold trials are rewarded with a bolus of water 50% of the time regardless of response. Clearly visible or invisible catch photic trials are used to ensure that the animal is responding correctly; rewarded is given only if the correct response is made. Catch stimuli are presented in 30% of trials. LabView (National Instruments, Austin, TX) is used to control task flow and monitor session progress. Visual stimuli are generated using a real-time visual stimulator (ViSaGe, Cambridge Research Systems, Rochester, Kent, England) and displayed on a CRT monitor (G90fb, ViewSonic, Walnut, CA). The NHPs are head-fixed using a minimally invasive technique and eye positions is tracked with an infrared camera (EyeLink 1000, SR Research, Mississauga, ON, Canada). Behavioral and neural data are collected using a Cerebus NeuroPort System (Blackrock Microsystems, Salt Lake City, UT, USA) at 10kHz. Data collection is conducted daily in 2 hour sessions.

### Photic Psychophysics

Pre-study values were collected during five task sessions conducted before implantation of the first array. Post-study values were collected across five days, 20 months following implant 1. This time period followed the implant and explant of array 1, implantation of array 2, and 16 months of stimulation. Pre and post study values were each averaged across five sessions, collected on consecutive day, with values weighted by session counts for each luminance.

During training, photic stimuli were presented randomly across the visual field space represented on the monitor, with a maximum eccentricity of 15° from center. Photic thresholds were calculated from a subset of presented stimuli localized to the area of array coverage, in polar coordinates from θ = 180 - 270°, and eccentricities of r = 1 - 14°. This range was determined from anatomical placement of the array combined with localization of receptive fields.

### Two-point discrimination task

This task was used to determine whether certain parameters evoke more than one percept as well as evaluate the distance at which stimulation on neighboring electrode produces separate percepts. The task will proceed with in a similar sequence to the threshold task previously described. The subject will place both hands on capacitive switches to indicate trial start. Following fixation within 1 degree of the fixation point displayed, either photic or electric stimuli will be presented. Photic stimuli will be Gaussian pulses presented on a CRT screen. Each trial presented either one or two pulses of light, the latter having a randomized variable distance between stimuli centers of 1.5 - 5 times the radius of the photic stimulus. Electric stimuli were presented simultaneously on pairs of two electrodes. All combinations of functional electrodes were tested to evaluate the range at which separate percepts can be evoked. One electrode pairing was conducted per session. The subject responded by lifting his left hand to indicate one stimulus observed and by lifting his right hand to indicate two observed percepts. Stimulation trials were randomly interdigitated with photic trial and accounted for 20-40% of trials within a session.

**Figure 9.**
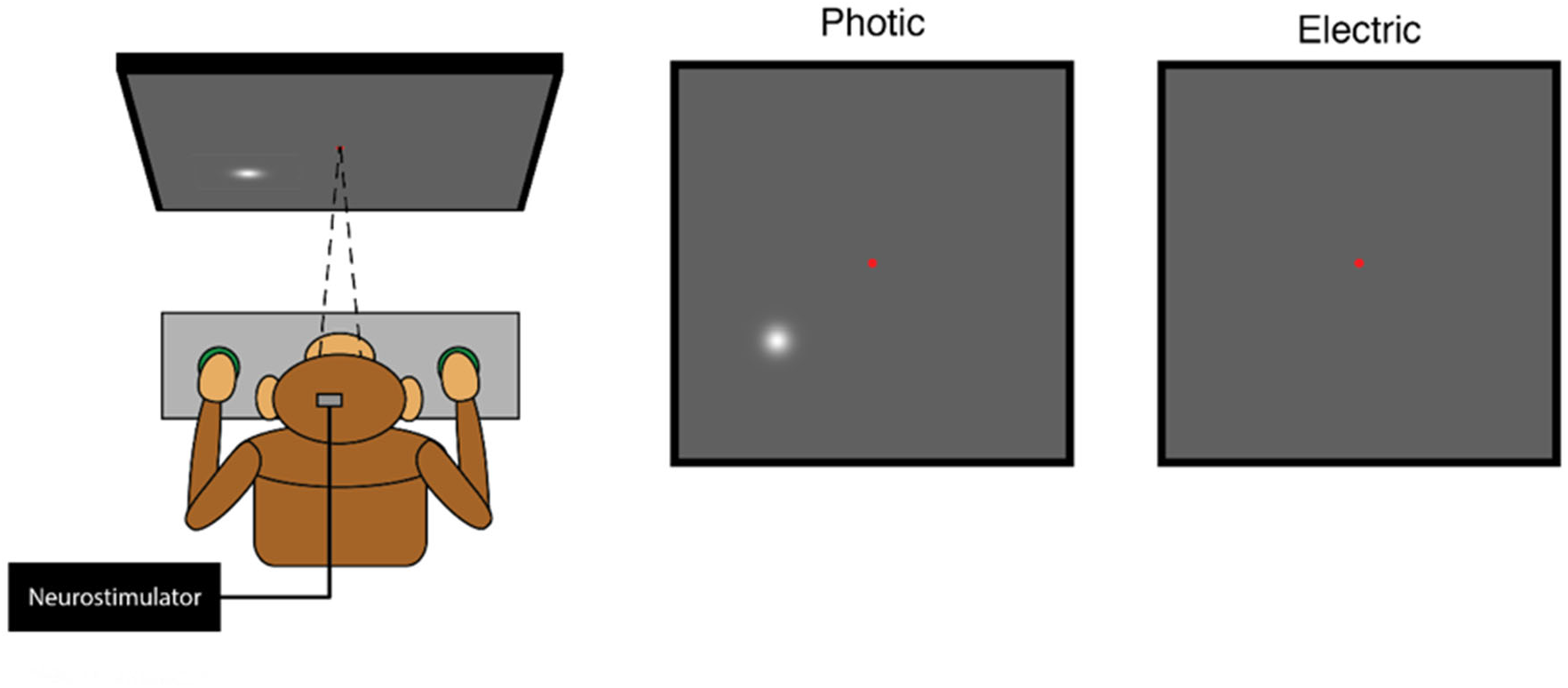
Task setup for all experiments. Non-human primate subjects are seated, head-fixed, facing a CRT monitor. Photic stimuli are present on monitor. Electric stimuli are delivered via an IZ2 neurostimulator with no photic stimuli presented on monitor, fixation point is maintained.

### Recording and stimulation

Recordings were conducted with a 128-channel Neural Signal processor (Blackrock Micro Systems). Stimulation experiments use an IZ2-128 stimulator (Tucker-Davis Technologies, Alachua, FL). The stimulator is capable of delivering 300μA to 128 channels. To achieve higher current levels, up to 3mA, a custom adder box was developed. Pulse trains consisted of symmetric, cathode-first, biphasic waveforms. The initial interphase interval and phase duration are set at 100μs and 600μs respectively, with pulse frequency and train duration set at 300Hz and 300ms.

### Data collection

Data was collected daily in sessions lasting up 3.5 hours. Sessions including electric stimulation were conducted most days, with gaps occurring for collection of receptive field data and photic behavioral data baselines.

### Determining electric thresholds

Electric stimuli were applied at 100μA increments from 0-900μA and consisted of10-40% of trials within a daily session, the reminder consisting of photic trials. At least one photic trial separated each electric trial to limit the effect of prior stimulations on the raw threshold collection. The amplitude of the current was applied randomly between the specified values. For each session, responses to stimulation were fit with a Weibull cumulative distribution function. The current value at 80% probability was determined to be the threshold value.

### Strength- and Charge-Duration Curves

Current thresholds for perception were collected for pulse duration from 0.1-0.7ms, for 300ms of 200Hz stimulation. Thresholds from these points were curve fit to Equation 1:

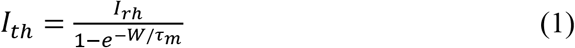

where *I_th_* is the threshold current, *I_rh_* is the rheobase current, *W* is the pulse width, and *τ_m_* is the membrane time constant. In order to curve fit, the rheobase current for a given parameter regime was set to the 10μA less than the minimum current threshold collected. The time constant was used to additionally tune the equation. The charge per phase threshold *Q_th_* was determined by the product of the current threshold and the pulse width, such that:

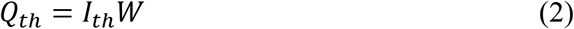

The minimum charge *Q_min_* was determined by the threshold charge at chronaxie.

### Predicting stimulation range and percept size

The radius of stimulation was determined using Equation 3.

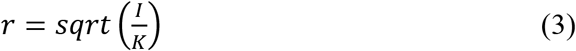

where *r* is the radius of stimulation for which tissue is directly activated, *I* is the applied current, and *K* is the current-distance constant. Calculations here use the value 675μA/mm^2^, based on V1 averaged values.

Phosphene size was predicted following a model developed based on data from surface stimulation of human visual cortex reported by [43].

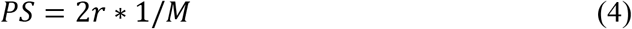

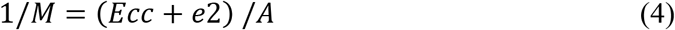

where *PS* is phosphene size, *M* is the linear cortical magnification factor, *Ecc* is the eccentricity, *e*2 is the eccentricity at which *M* is half of its foveal value, and *A* is a cortical scaling factor. Based on the reported values, this calculation used *e*2 = 3.67 *°*. Since the study reported a range of *A* from 15 to 45, these values were used as bounds to create a range of expected percept sizes based on the upper and lower bounds.

## Acknowledgements

This work was funded by the Medical Technology Enterprise Consortium: Project Call, MTEC-16-02-BMl-06; Maximizing Vision Restoration through Optimizing Brain Computer Interface Design, Base Agreement, 2017-608, Award Number, 001. Contract Number: W81XWH-15-9-0001.

